# Stochastic competitive exclusion leads to a cascade of species extinctions

**DOI:** 10.1101/069203

**Authors:** José A Capitán, Sara Cuenda, David Alonso

## Abstract

Community ecology has traditionally relied on the competitive exclusion principle, a piece of common wisdom in conceptual frameworks developed to describe species assemblages. Key concepts in community ecology, such as limiting similarity and niche partitioning, are based on competitive exclusion. However, this classical paradigm in ecology relies on implications derived from simple, deterministic models. Here we show how the predictions of a symmetric, deterministic model about the way extinctions proceed can be utterly different from the results derived from the same model when ecological drift (demographic stochasticity) is explicitly considered. Using analytical approximations to the steady-state conditional probabilities for assemblages with two and three species, we demonstrate that stochastic competitive exclusion leads to a cascade of extinctions, whereas the symmetric, deterministic model predicts a multiple collapse of species. To test the robustness of our results, we have studied the effect of environmental stochasticity and relaxed the species symmetry assumption. Our conclusions highlight the crucial role of stochasticity when deriving reliable theoretical predictions for species community assembly.

## 1. Introduction

Ecological communities are shaped from the complex interplay of four fundamental processes (Vellend, 2010): selection, in the form of species interactions that favor certain species against others; speciation, leading to the appearance of new species, better adapted to the environment; dispersal, which permits spatial propagation of individuals; and ecological drift, a demographic variability in species population numbers due to the stochastic processes that take place. Ecological drift, in particular, has a prevalent role in modern theoretical frameworks in community ecology (Black and McKane, 2012). Accordingly, current approaches reveal the need for process-based, stochastic models that help to understand how ecological communities are assembled and their interaction with environmental factors (Wisz et al., 2013).

Classical community ecology, however, has mainly relied on deterministic community models (see Roughgarden (1979) and references therein), most of them based on Lotka-Volterra dynamics, although alternatives have been proposed (Schoener, 1974a). There is a long-standing research focus on community assembly models, in which communities are built up through species invasions, and most of them rely on deterministic approaches (Post and Pimm, 1983; Law and Morton, 1993, 1996; Capitán et al., 2009; Capitán and Cuesta, 2011; Capitán et al., 2011). On the other side, there have been strong theoretical efforts to describe community assemblages in stochastic terms (Hubbell, 2001; Alonso et al., 2008; Rosindell et al., 2011). In certain situations, the results and conclusions derived from deterministic models have been shown to be quite different in the presence of stochasticity (Bolker and Grenfell, 1995; Alonso et al., 2007; Haegeman and Loreau, 2011; Bonachela et al., 2012; Wang et al., 2012).

One of the contexts where the differences between deterministic and stochastic approaches become apparent is related to theoretical formulations of the competitive exclusion principle (Volterra, 1926; Gause, 1934; Hardin, 1960). This principle constitutes a fundamental pillar of community ecology and belongs to the traditional body of ecological theory. It provides a useful theoretical framework to explore how complex species assemblages persist over time. Important concepts such as adaptation to shared niches (Roughgarden, 1979), species limiting similarity (MacArthur and Levins, 1967; Roughgarden, 1974) or niche partitioning (Pielou, 1977; Schoener, 1974b) all are immediate derivations of the principle. Classical approaches predict the maximum degree of species similarity that permit species stable coexistence (MacArthur, 1969, 1970). However, theoretical predictions for limiting similarity often rely on deterministic community models (see MacArthur (1968); Levin (1970); Haigh and Maynard-Smith (1972); Chesson (1990) and Appendix A for a discussion on competitive exclusion based on deterministic approaches), and the relevance of stochasticity, in the form of ecological drift, to species coexistence has remained almost unexplored (with the exception of Turelli (1980)). The relationship between limiting similarity and environmental stochasticity has been studied more thoroughly (May and MacArthur, 1972; Turelli, 1978, 1981).

Recently, we focused on the influence of ecological drift on the similarity of coexisting species via the competitive exclusion principle (Capitán et al., 2015). In that contribution we showed that, in the presence of ecological drift, the maximum degree of similarity that ensures stable coexistence can be significantly lowered when compared to the corresponding limits to similarity derived from deterministic models. If similarity is interpreted in terms of an interspecific competitive overlap (MacArthur and Levins, 1967; Roughgarden, 1974), stochasticity displaces the deterministic threshold towards lower values of the competitive overlap (Capitán et al., 2015). Thus, when stochasticity is considered, the extinction phenomena caused by competitive exclusion takes place at lower values of the competitive overlap (i.e., species have to be more dissimilar to stably coexist in the presence of ecological drift).

Ecological drift becomes a key process determining species coexistence in aspects other than the maximum similarity of co-occurring species. Beyond a more restrictive threshold in competition induced by ecological drift (which was the main result of Capitán et al. (2015)), we here analyze the influence of demographic stochasticity on the extinction mechanism itself, which in principle can lead to either sequential or grouped extinctions as competition strength increases. For that purpose, we considered a deterministic, Lotka-Volterra model and its stochastic counterpart, both of which treat species interactions symmetrically. Whereas the deterministic model predicts the multiple extinction of all the species in the community but one as competition crosses over a certain threshold, in the presence of demographic stochasticity extinctions proceed progressively, in the form of a cascade, as competition increases. The only difference between both approaches is the explicit consideration of ecological drift in the dynamics. In order to derive our conclusions, we developed convenient analytical approximations to the steady-state configurations of the stochastic system for simple species assemblages formed by two or three species. Such approximations help us to partition the set of feasible population numbers into regions associated to coexistence, or the extinction of one, two, or three species. The steady-state probabilities, when aggregated over those regions, unveil the extinction cascade phenomenon. Our main result reveals overlapping windows in competitive strength, at low values related to configurations where the coexistence of three species is the most probable state, intermediate ranges where it is more likely to observe two-species assemblages, and large competition values for which the most probable state is formed by one species or none. We also studied the transition to the deterministic model when demographic stochasticity tends to zero, and our results reveal an abrupt transition to situations compatible with small stochasticity.

To test the robustness of our conclusions, we replaced demographic stochasticity by environmental stochasticity and confirmed that, although the extinction phenomena are qualitatively different, the extinction cascade persists. We also relaxed the assumption of symmetry to assess the effect of stochasticity on deterministic models that not only predict multiple extinctions, as in the fully symmetric scenario, but also lead to both progressive and grouped extinctions for fixed competitive strengths. When stochasticity comes into play, however, the stochastic cascade persists and the expected extinction sequence is qualitatively different from its deterministic counterpart. Thus, the predictions of both models are significantly different in generic, non-symmetric scenarios for species interactions.

The paper is organized as follows: in Section 2 we describe both the deterministic and the stochastic frameworks, the latter based on the formulation of Haegeman and Loreau (2011), and show that the deterministic, symmetric model predicts a multiple species extinction. In Section 3 we start by presenting the analytical approximations for a two-species stochastic community model, and we then extend the procedure to a three-species community. These approximations help us to obtain analytical formulae for the critical points of the steady-state, joint probability distribution of the community. Formulae for saddle points are then used to properly define aggregated probabilities of coexistence, or one-, two-, and three-species extinction, which reveal themselves the sequential decline of species driven by ecological drift. After studying the small stochasticity limit and testing the robustness of our results, we conclude the paper with several implications and prospects (Section 4).

## 2. Model description

For the sake of simplicity, in this contribution we will focus on the symmetric version of the deterministic Lotka-Volterra competitive dynamics (see Appendix A),

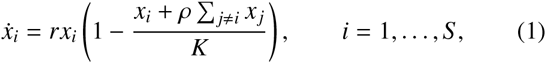

where *x_i_* stands for the population density of species *i* (space is implicitly assumed) and model parameters are uniform and species-independent. Here *r* stands for an intrinsic, species-independent growth rate, *ρ* measures interspecific competition, *K* represents a carrying capacity, and *S* is the species richness of the community. The dynamics has an interior equilibrium point, 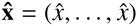, where 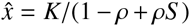, which is globally stable if and only if *ρ* < 1 (Hofbauer and Sigmund, 1998; Capitán et al., 2015). In the symmetric scenario, the competitive exclusion principle adopts a very simple formulation (see Appendix A for further details on the general, non-symmetric case). A complete stability analysis of the boundary equilibrium points shows that, for *ρ* > 1, all the species become extinct except for just one of them (see Appendix B). As a consequence, competitive exclusion in the symmetric, deterministic model implies the joint extinction of *S* – 1 species.

We now explicitly incorporate ecological drift (demographic stochasticity) in the symmetric scenario in order to show that species are sequentially displaced in the presence of stochasticity due to competitive exclusion, following a cascade of extinctions, as competition strengthens. A standard way to extend deterministic models to incorporate ecological drift is deeply described in Haegeman and Loreau (2011). The state of the system is described by the vector of population numbers *n_i_* at time *t*, **n**(*t*) = (*n*_1_(*t*),…, *n_S_* (*t*)). Contrary to the deterministic case, which focuses on population densities *x_i_* = *n_i_*/Ω, Ω being a meaningful measure (area, volume) of the system size, here discrete population numbers are considered. The elementary processes that define the stochastic dynamics (local births and deaths, immigration, and competition) are characterized by probability rates that, in the deterministic limit, yield the Lotka-Volterra equations (1). As in Haegeman and Loreau (2011), we choose the following probability rates to model elementary processes:

1. Local births (deaths) of species *i* occur at a density-independent rate *r*^+^ *n_i_* (*r*^−^*n_i_*). We adopt the notation *r* = *r*^+^ – *r*^−^ to represent the net growth rate in the absence of competitors.
2. Immigration of a new individual of species *i* takes place at a rate *μ*. Although the deterministic model (1) does not include immigration, dispersal is an important process driving community assembly (Vellend, 2010). In addition, immigration is key for the stochastic process to reach a nontrivial steady-state. We consider here the low-immigration regime, in which the deterministic limit is expected to recover results close to those yielded by Eq. (1), see Capitán etal. (2015).
3. Intraspecific competition occurs at a density-dependent rate 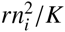, where *K* represents a species carrying capacity.
4. Interspecific competition between species *i* and *j* (*i* ≠ *j*) takes place at a probability *rρn_i_n_j_*/*K* per unit time (it is also a density-dependent rate).

Population vectors **n**(*t*) belong to the configuration space 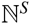, where 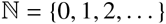. The elementary processes listed above define a birth-death-immigration stochastic process in continuous time, and the probability *P*(**n**, *t*) of finding a population vector **n**(*t*) at time t is determined by the master equation,

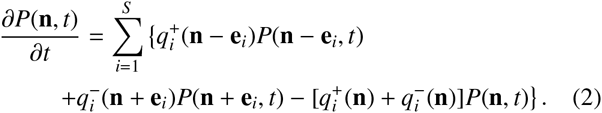

Here 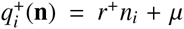 is the overall birth rate for species *¡*, 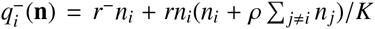 is the overall death rate, and **e**_*i*_ = (0,…, 1,…, 0) is the *i*-th vector of the canonical basis of 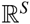. Notice the correspondence of these rates with the terms arising in the deterministic model (1) for *μ* = 0. The steady-state probability distribution is obtained by solving the coupled recurrence equation given by the condition *∂P*(**n**, *t*)/*∂t* = 0 (see Appendix C for details).

Our approach develops analytical approximations for the critical points of the joint probability function, not for the probability itself. For *S* = 1 the stationary state of the birth-death-immigration model can be exactly solved in terms of hypergeometric functions (Haegeman and Loreau, 2011). For *S* > 1, in the absence of competition (*ρ* = 0) populations are uncorrelated, and the joint probability distribution factors as a product of marginal probabilities, which reduces this case to a onedimensional problem. The fully neutral model (*ρ* = 1)for *S* > 1 can be solved as well (Haegeman and Loreau, 2011). We cannot find analytically the steady-state distribution for *S* ≥ 2 and *ρ* > 0, though. Haegeman and Loreau (2011) devised approximations for this case, but we will follow here a different approach to find analytical formulae for the critical points of the steady-state distribution in the case of small-sized communities.

## 3. Results

In Capitán et al. (2015) we showed numerically that the steady-state distribution for two-species communities presents a maximum at an interior point of the configuration space, as well as two boundary maxima with population vectors of the form (*n*, 0) and (0, *n*). This implies that the discrete probability distribution, when extended to be real-valued (by, for instance, cubic spline interpolation), must exhibit by continuity two saddle points located in between the coexistence maximum and the two boundary maxima. These saddle points can be used to conveniently partition the configuration space into regions associated to coexistence, the extinction of one species, or the extinction of two species. In this section we develop analytical approximations that help us to obtain good estimates for the critical points of the joint probability distribution. We use the *S* = 2 case to illustrate the technique. We then extend the method for three-species assemblages, and use the approximated saddle points to partition the configuration space into regions for coexistence and for the three possible states where extinctions have occurred. The aggregation of the joint probability over those regions unveils the extinction cascade phenomenon.

### 3.1. Critical points for two-species communities

To estimate the location of the critical points of the joint distribution *P*(*n*_1_, *n*_2_) we use that the conditions *∂P*(*n*_1_, *n*_2_)/*∂n*_1_ = 0 and *∂P*(*n*_1_, *n*_2_)/*∂n*_2_ = 0 are equivalent to

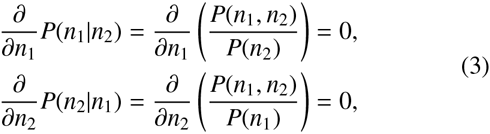

*P*(*n*_1_|*n*_2_) being the probability that species 1 has *n*_1_ individuals conditioned to species 2 having *n*_2_ individuals. This means that critical points of the joint distribution *P*(*n*_1_, *n*_2_) can also be obtained through conditional probabilities. By fixing *n*_2_, we just need to evaluate the derivative of *P*(*n*_1_|*n*_2_) along the *n*_1_ direction. The same applies under the change *n*_1_ ↔ *n*_2_.

#### 3.1.1. Approximated conditional probabilities

We now approximate *P*(*n*_2_|*n*_1_) by *T*(*n*_2_|*n*_1_) as follows. For *S* = 2, the steady-state distribution satisfies a two-term recurrence in population numbers *n*_1_ and *n*_2_, namely

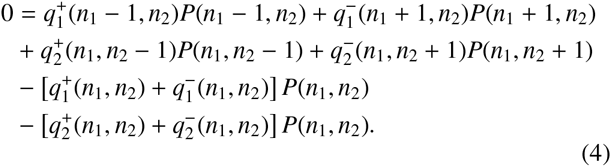

Notice that we are interested in approximating the conditional probability *P*(*n*_2_|*n*_1_) where *n*_1_ is fixed. Hence, in the approximation we ignore the terms in Eq. (4) that involve variation of *n*_1_, and assume that *n*_1_ acts as a fixed parameter in the remaining terms. Thus, the approximated conditional probability *T*(*n*_2_|*n*_1_) satisfies

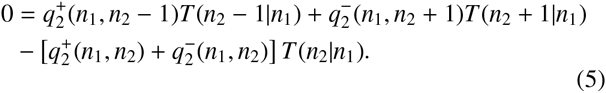

This expression fulfills a detailed balance condition (Karlin and Taylor, 1975), which yields an approximate one-term recurrence formula in *n*_2_,

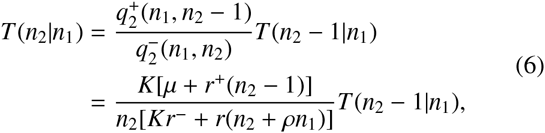

and leads to an explicit solution in terms of hypergeometric functions, as in Haegeman and Loreau (2011). A symmetric recurrence holds for *T*(*n*_1_|*n*_2_).

#### 3.1.2. Analytical formulae for critical points

Our next step is to approximate the partial derivative along the n1 direction by the backwards discrete difference (comparable results are obtained with the forward difference),

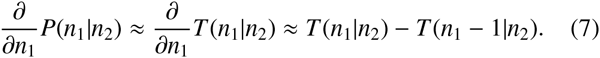

Similarly,

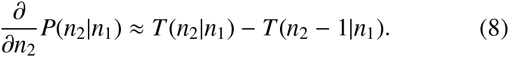

Eq. (6) allows us to write the system for the critical points of the two-dimensional joint-probability surface, *∂P*(*n*_1_|*n*_2_)/*∂n*_1_ = *∂P*(*n*_2_|*n*_1_)/*∂n*_2_ = 0, as

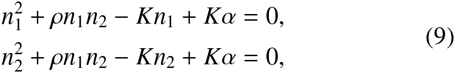

where *α* = (*r*^+^ – *μ*)/*r*. Solving the quadratic system yields the following estimates for the interior critical points: *M*_1_ = (*m*_+_, *m*_+_), which can be either a local maximum or a saddle point depending on the value of *ρ* (see below), and the local minimum *M*_2_ = (*m*_−_, *m*_−_), where

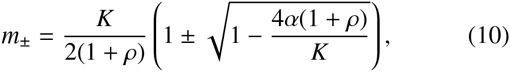

and two (symmetrical) saddle points *Q*_1_ = (*s*_+_, *s*_−_) and *Q*_2_ = (*s*_−_, *s*_+_), where

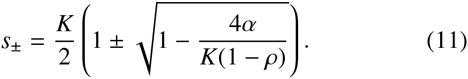

To test the goodness of these analytical approximations, critical points of the exact probability distribution are determined numerically by extending the discrete distribution *P*(*n*_1_, *n*_2_) to be a real-valued function using cubic spline interpolation both in the *n*_1_ and *n*_2_ directions. We then solve numerically the system *∂P*(*n*_1_, *n*_2_)/*∂n*_1_ = *∂P*(*n*_1_, *n*_2_)/*∂n*_2_ = 0, using analytical predictions as initial guesses for iterative root finding. The Hessian matrix decides whether a given point is a local maximum, a local minimum or a saddle point. We find that the critical points are in excellent agreement with the analytical formulae above (see Fig. 1a, in which we plot the coordinates of each critical point calculated both analytically and numerically).

**Figure 1:**
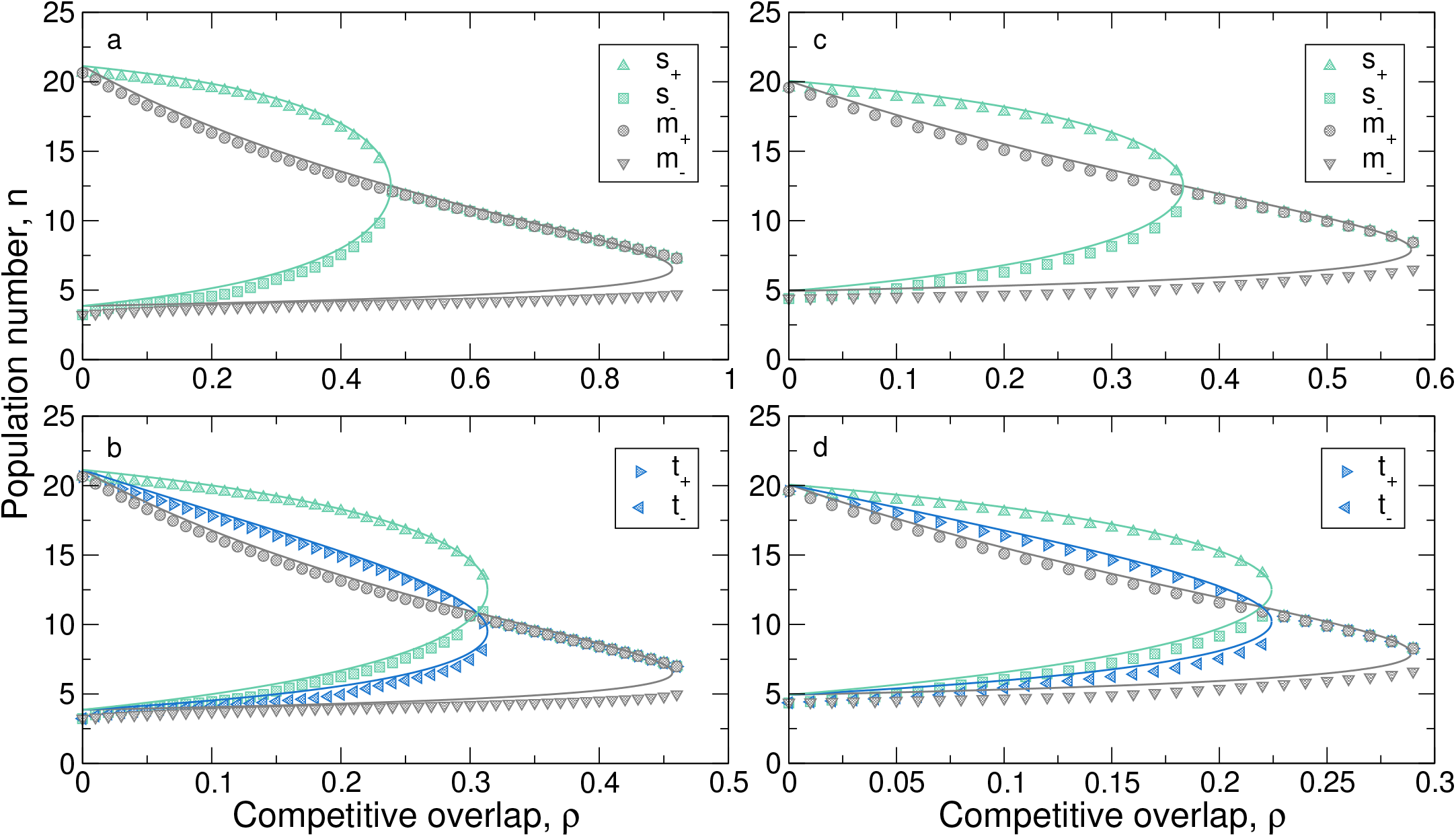
(a) Coordinates of the critical points for *S* = 2 as a function of *ρ*. They were calculated using the exact steady-state distribution (symbols) as well as the approximations given by Eqs. (10) and (11) (lines). Once the coexistence maximum *M*_1_ = (*m*_+_, *m*_+_) and the two saddle points *Q_i_* = (*s*_±_, *s*_+_) (*i* = 1,2) coalesce, *M*_1_ becomes a saddle point (there is no longer a coexistence maximum). Model parameters are *r*^+^ = 50, *r*^−^ = 35, *μ* = 1, *K* = 25. (b) Coordinates of the critical points for a three-species community (symbols), compared with theoretical approximations (17) and (19). When the two saddle points coincide, the former maximum becomes a saddle point. Parameter values are the same as in panel (a). Panels (c)-(d) check the accuracy of our approximations for other set of parameter values, namely *r*^+^ = 10, *r*^−^ = 7.5, *μ* = 0.1, and *K* = 25. As before, (c) represents the coordinates of the critical points for *S* = 2 and (d) for *S* = 3.

For *ρ* < *ρ_c_*, with

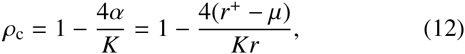

the analysis of the Hessian matrix reveals that *M*_1_ is a (coexistence) local maximum. For *ρ* = *ρ_c_*, saddle points *Q*_1_ and *Q*_2_ coincide with *M*_1_ and, for *ρ* > *ρ_c_*, *M*_1_ transforms into a saddle point and the former saddle points *Q*_1_ and *Q*_2_ (cf. Eq. (11)) no longer exist. Moreover, when *ρ* > *K*/4*ρ* – 1, all interior critical points (10) become complex and the only persistent maxima are those located at the boundary.

There are four critical points at the boundary, which can be approximated using Eq. (6) for *n*_1_ = 0, resulting in two local maxima [(*b*_+_, 0), (0, *b*_+_)] and two local minima [(*b*_−_, 0), (0, *b*_−_)], where

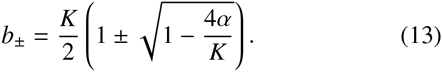

Observe that the non-zero coordinates of the local maxima (minima) coincide with those of the interior maximum (minimum) for *ρ* = 0.

### 3.2. Critical points for three-species communities

The method proposed in the previous subsection can be fully extended to the case of a three-species community. Critical points are obtained by approximating conditional probabilities *P*(*n*_1_|*n*_2_, *n*_3_) and taking discrete derivatives with respect to the first argument. As before, we consider the three-term recurrence relation that fulfills the joint distribution *P*(*n*_1_, *n*_2_, *n*_3_) and ignore the terms that involve variation in population numbers *n*_2_ and *n*_3_. Under this approximation, the steady-state condition reduces to

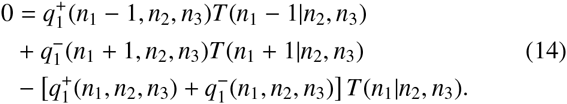

Due to detailed balance, the approximate conditional probabilities *T*(*n*_1_|*n*_2_, *n*_3_) satisfy the one-term recurrence relation

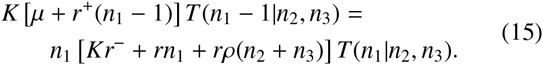

Repeating for *S* = 3 the procedure devised to estimate the coordinates of the critical points leads to the set of quadratic equations

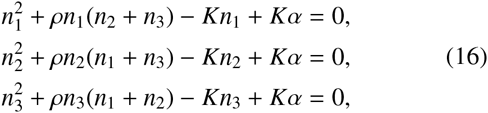

which yields 8 interior critical points, 6 of which are saddle points, and the remaining two points are a local minimum and, as in *S* = 2, a point that is a maximum or a saddle point depending on p. Explicit expressions for their coordinates are given below. For the sake of comparison we have also calculated critical points using the exact joint distribution *P*(*n*_1_, *n*_2_, *n*_3_), see Fig. 1b.

The coordinates for the interior critical points are: on the one hand, *M*_1_ = (*m*_+_, *m*_+_, *m*_+_) and *M*_2_ = (*m*_−_, *m*_−_, *m*_−_), where

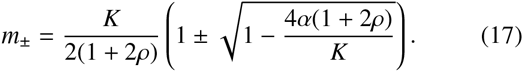

Both solutions turn out to be complex when

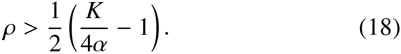

On the other hand, six interior critical points appear at points *Q_i_*, *i* = 1,…, 6, where *Q*_1_ = (*t*_+_, *t*_+_, *s*_−_) and *Q*_2_, *Q*_3_ are obtained as the cyclic permutations of the entries of *Q*_1_, whereas *Q*_4_ = (*t*_−_, *t*_−_, *s*_+_) and the entries of *Q*_5_ and *Q*_6_ are the cyclic permutations of that of *Q*_4_, with

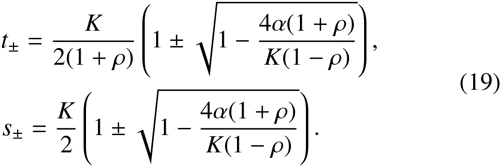

Both *s*_±_ and *t*_±_ are real whenever *ρ* < *ρ_c_*, where

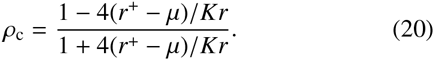

In spite that Eqs. (17) and (19) have been obtained using an approximate form for conditional probabilities, the numerical calculation of the critical points using the actual distribution is in very good agreement with these approximations (see Fig. 1b). For *ρ* < *ρ*_c_, *M*_1_ is classified as a local maximum. At *ρ* = *ρ*_c_ all saddle points and *M*_1_ coalesce in a single point. For *ρ* > *ρ*_c_, however, *M*_1_ becomes a saddle point, as can be checked numerically with the Hessian matrix.

Boundary maxima are of the form (*n, n*, 0), (*n*, 0, 0)—and their cyclic permutations. The non-zero coordinates of the former are equal to that of the coexistence maximum *M*_1_ obtained for 2 species, see Eq. (10); the non-zero entries of the latter are the same as the boundary maxima for *S* = 2, see Eq. (13).

#### 3.2.1. Configuration-state partitioning

For *S* = 2 potential species, a simple way to divide the configuration space is to use saddle points. Fig. 2 depicts steady-state distributions for increasing competitive overlap as well as the location of saddle points. A natural partitioning should relate configurations around the coexistence maximum to species coexistence, and states near the boundary maxima to configurations close to one-species extinction. As Fig. 2 shows, saddle points discriminate with accuracy the configurations that can be associated to coexistence from those that can be related to one-species extinction. According to the coordinate (*s*___) that closest to the boundary (cf. Eq. (11)), the partitioning results as:

1. 0 ≤ *n*_i_ < *s*_−_ for *i* = 1,2. This square is associated with full extinction.
2. 0 ≤ *n*_1_ < *s*_−_, *n*_2_ > *s*_−_ or 0 ≤ *n*_2_ < *s*_−_, *n*_1_ > *s*_−_. These two rectangles are associated with the extinction of one species, since configurations are close to extinction in the form (*n*, 0) or (0, *n*).
3. *n_i_* > *s*_−_ for *i* = 1,2. The rest of the configuration space puts together states that can be associated to coexistence.

**Figure 2:**
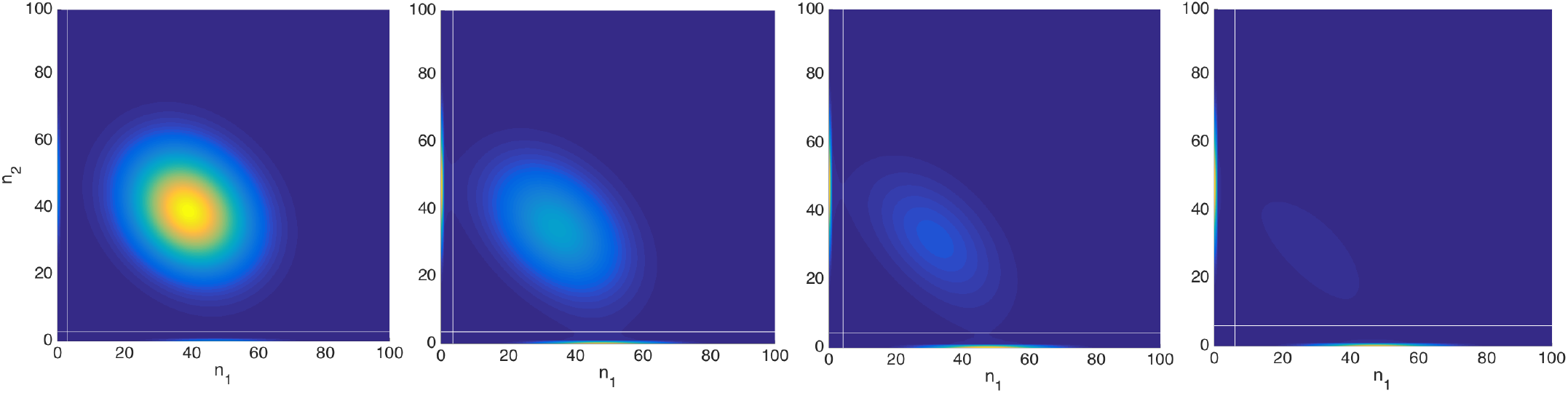
Partitioning of the configuration space 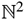 for *S* = 2 species for (from left to right) *ρ* = 0.2, *ρ* = 0.35, *ρ* = 0.45 and *ρ* = 0.6; remaining parameters are *r*^+^ = 50, *r*^−^ = 27, *μ* = 1, and *K* = 50. The smaller coordinates of the two saddle points (used to draw white lines) are used to partition the space into regions associated to coexistence (central square), one-species extinction (the two rectangles which contain the boundary maxima) and two-species extinction (the small square that contains the origin). As competition increases, the coexistence maximum approaches to the origin, and saddle points become closer to the maximum.

Note that, strictly speaking, there will be configurations where both *n*_1_, *n*_2_ > 0 are classified as one- or two-species extinction states. The classification here is meant to separate configurations that are close to boundary maxima in which one or two species are extinct from those that can be associated to coexistence, in which species populations are far from being extinct.

The partitioning for three-species communities simply generalizes the *S* = 2 case. Again, each saddle point has a coordinate close to the boundary (cf. *s*_−_ and *t*_−_ in Eq. (19)). Note also that *s*_−_ > *t*_−_. Taking this fact into account, saddle points divide the configuration space 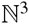 into several regions, which have been depicted in Fig. 3:

1. 0 ≤ *n*_i_ < *t*_−_ for *i* = 1,2,3. This cube is associated with the extinction of the three species (since the origin belongs to this region).
2. 0 ≤ *n_i_* < *t*_−_ for *i* = 1,2 and *n*_3_ > *t*_−_ (and the two remaining combinations). These three parallelepipeds are associated with the extinction of two species, because boundary maxima of the form (*n*, 0, 0)—and its cyclic permutations— are situated inside this volume, as well as other population configurations close to two-species extinctions.
3. *n_i_* > *s*_−_ for *i* = 1, 2, 3. This cube contains the interior maximum and is therefore associated with the coexistence of the three species.
4. The remaining volume of the configuration space, where boundary maxima of the form (*n, n*, 0)—and its cyclic permutations—are located, is associated with the extinction of one species.

**Figure 3:**
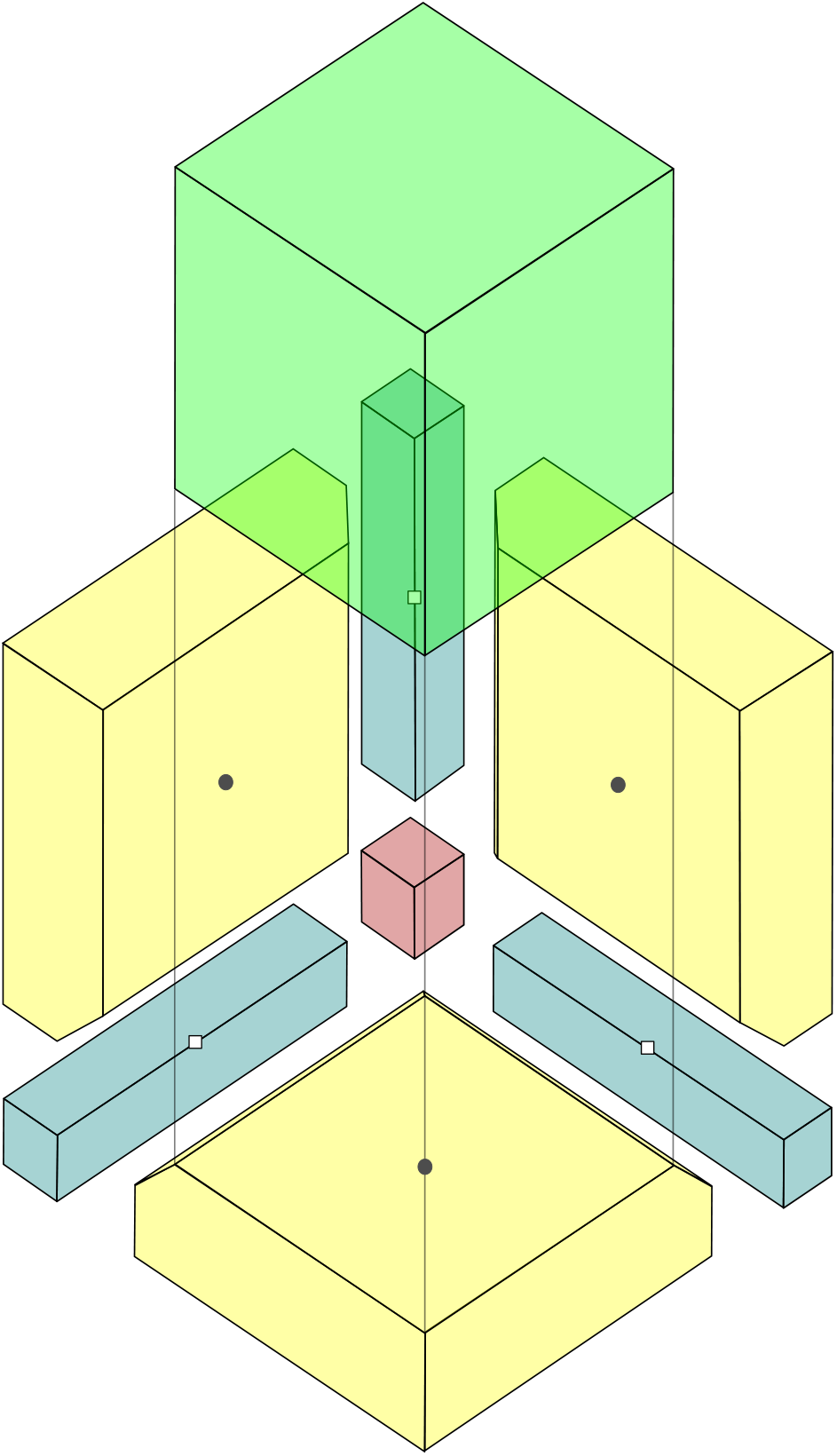
Partition of the configuration space 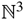 for *S* = 3 potential species. To ease visualization, we have separated 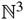 into four regions according to saddle points: black circles denote points *Q*_1_, *Q*_2_ and *Q*_3_, whereas white squares represent saddle points *Q*_4_, *Q*_5_ and *Q*_6_. The latter are used to define regions for complete (red) and two-species extinctions (blue), and the former determine the coexistence volume (green) and the one-species extinction region (yellow). The (green) cube has been displaced vertically to facilitate visualization.

The configuration-state partitioning slightly differs from the general case when saddle points have coalesced. If *ρ* > *ρ*_c_, the point *M*_1_ = (*m*_+_, *m*_+_, *m*_+_) is classified as the only saddle point. Based on its coordinates, the configuration space is partitioned as follows:

1. 0 ≤ *n_i_* < *m*_+_ for *i* = 1,2,3. This cube is associated with full extinction.
2. 0 ≤ *n_i_* < *m*_+_ for *i* = 1, 2 and *n*_3_ > *m*_+_ (and the two remaining combinations). These three parallelepipeds are associated with the extinction of two species.
3. *n_i_* > *m*_+_ for *i* = 1, 2, 3. This cube is associated with coexistence configurations.
4. The remaining volume of the configuration space is related to the extinction of one species.

The same partitioning applies for the *S* = 2 case when only a single saddle point remains.

### 3.3. Extinction cascade

Saddle points of the joint probability distribution have allowed us to establish a natural partitioning into regions associated to coexistence (containing the coexistence maximum) and to the extinction on one, two, or three species (containing the corresponding boundary maxima), see Fig. 3. Over these regions we aggregate the joint distribution *P*(*n*_1_, *n*_2_, *n*_3_), calculated numerically as described in Appendix C, to define the overall probability of coexistence,

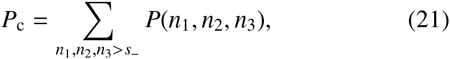

the probability of three-extinct species configurations,

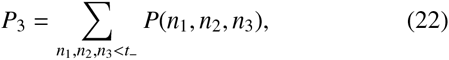

and the probability of two-extinct species, which by symmetry over cyclic permutations of (*n*_1_, *n*_2_, *n*_3_) can be expressed as

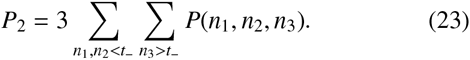

The probability of one-extinct species, P1, is obtained from the normalization condition *P*_c_ + *P*_1_ + *P*_2_ + *P*_3_ = 1. Fig. 4 shows these aggregated probabilities as a function of *ρ* for two sets of model parameters. In the first case, the coexistence probability is almost one for low values of the competitive overlap and, as *ρ* increases, at some point the probability declines rapidly. At the same time, the probabilities of one and two extinct species begin to increase. Note that, once the threshold pc has been crossed over, the most probable state consists of a single, extant species, and the probability of coexistence becomes negligible.

**Figure 4:**
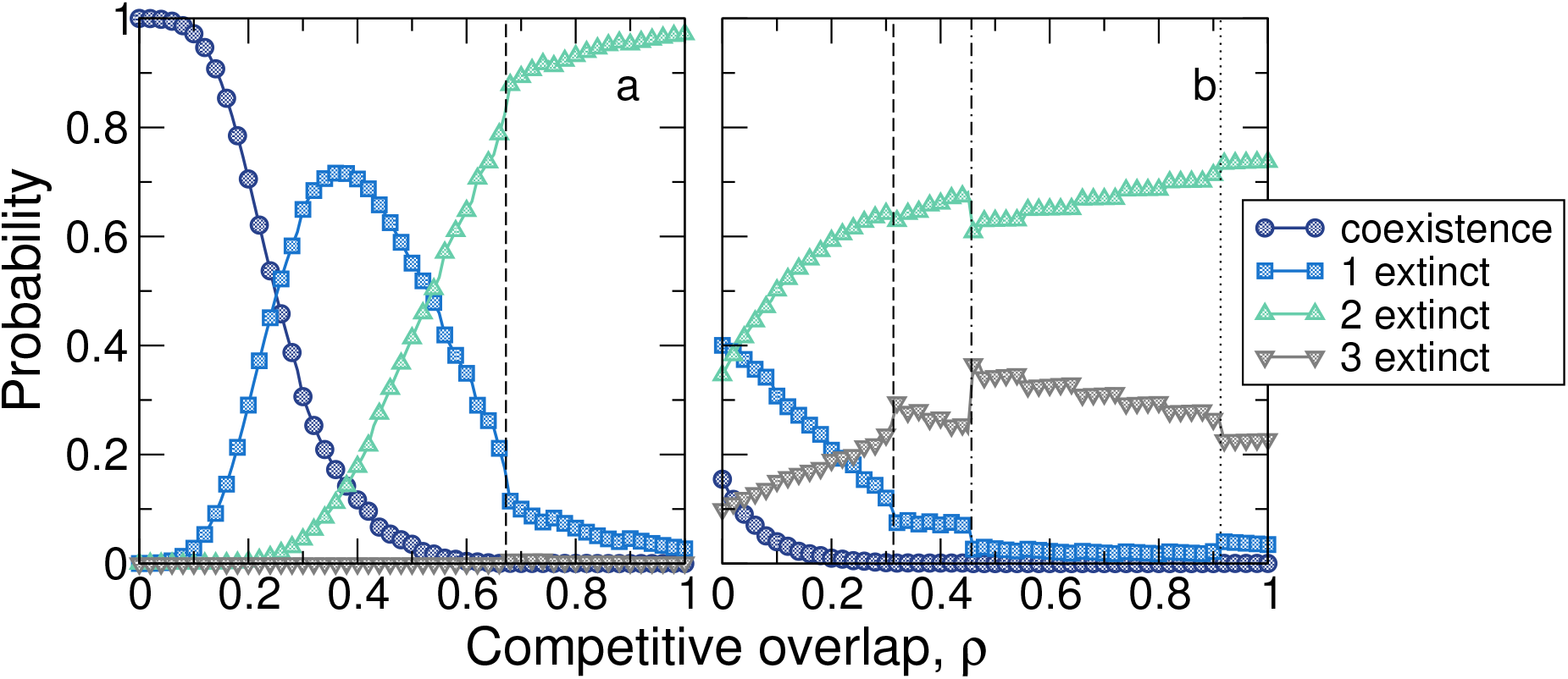
(a) Stochastic extinction cascade. The panel shows the dependence of aggregated probabilities over the regions determined by the critical points as a function of competitive overlap *ρ*. Parameter values are *S* = 3, *r*^+^ = 50, *r*^−^ = 10, *K* = 25, and *μ* = 1. Although initially the three species coexist, at intermediate values of *ρ* the most probable state is formed by only two extant species. For large competition values the most likely configuration comprises a single extant species. A vertical, dashed line marks the value *ρ*_c_ (cf. Eq. (20)) at which the probability of coexistence is negligible. (b) Same as panel a but for a higher intrinsic mortality rate, *r*^−^ = 35. In this case, the aggregated probability of complete extinction is non-negligible even for *ρ* = 0. Aggregated probability for one-extinct species configurations starts declining and, at the same time, the two-extinct species configurations become more likely as *ρ* increases. Close to *ρ* = 1 the system alternates most of the time between a single-species state or a completely extinct community. The vertical, dashed line marks the threshold *ρ*_c_. The dot-dashed line shows the value given by Eq. (18), at which the two critical points *M*_1_ and *M*_2_ no longer exist. Finally, a dotted, vertical line marks the value (*ρ* = *K*/4*α* – 1, see Section 3.1) at which the boundary maxima of the form (*n, n*, 0)—and permutations—no longer exist.

In the second case, corresponding to a larger value of the mortality rate *r*^−^, the threshold *ρ*_c_ at which the coexistence maximum *M*_1_ transforms into a saddle point becomes smaller. The probability of coexistence rapidly declines as *ρ* increases and, in addition, there is a non-negligible probability of complete extinction. Remarkably, Fig. 4b shows that, for smaller values of the carrying capacity, coexistence is not the only possible state even at *ρ* = 0. This puts a practical limit to the maximum number of coexisting species which does have a deterministic counterpart—recall that, due to global stability, the deterministic model permits the packing of an arbitrary number of species for *ρ* < 1.

Contrary to the deterministic prediction that *S* – 1 extinctions take place abruptly as *ρ* increases (Appendix B), Fig. 4 shows that ecological drift induces a sequential cascade of extinctions, in which states with a larger number of extinct species are more prone to be observed as the competitive overlap increases.

An important remark is on purpose here. The cascade of extinction we have just described has nothing to do with the degree of synchronicity in which extinctions take place along time, i.e., the term “cascade” does not refer here to a sequential extinction in time. In particular, the symmetric, deterministic model leads in general to asynchronous extinctions. The stochastic phenomenon analyzed in this contribution refers to the progressive extinctions that occur as competitive strength increases.

### 3.4. Limit of small demographic stochasticity

In the absence of stochasticity, the deterministic model predicts the extinction of *S* – 1 species once the threshold in competition *ρ* = 1 is crossed over. In the stochastic case, the extinction threshold is pushed to smaller values of competitive strength (Capitán et al., 2015), and the probabilities of configurations with one or more extinct species are non-zero in overlapping windows of competition. These two scenarios only differ on the presence or absence of demographic stochasticity, but lead to significantly different outcomes. Therefore, incorporating demographic stochasticity appears to be very relevant in the dynamics of ecological community models.

In order to evaluate the importance of demographic stochasticity, we have tried to quantify the difference between these two scenarios as stochasticity decreases. To do so, we have studied the limit of small stochasticity, in which the deterministic model is to be recovered. As shown below, the transition to the small-stochasticity scenario is abrupt, hence incorporating demographic stochasticity to community models should be strongly considered.

Since fluctuations are expected to decrease as population size increases (see Appendix D), the small-stochasticity limit is equivalent to the limit of large population sizes, so we have repeated the analysis by increasing the carrying capacity at fixed *ρ*. We have quantified the intensity of demographic noise by the coefficient of variation of population abundances, *v* = *σ_n_*/〈*n*〉, *σ_n_* being the standard deviation of population numbers and 〈*n*〉 the average value. As shown in Appendix D, when *K* ≫ 1 then *σ_n_* ~ *K*^1/2^ and 〈*n*〉 ~ *K*, so *v* tends to zero in the limit of large population sizes. From the numerical point of view, to get close to the deterministic scenario we would have to choose a carrying capacity value such that the average 〈*n*〉 is large enough compared to the variability in populations. In practice, we will assume that the system is close to a low-noise regime when the actual coefficient of variation, obtained though the joint probability distribution of the stochastic model calculated numerically, is close to that obtained by a Gaussian approximation of the joint distribution valid in the limit *K* ≫ 1 (see Appendix D). Note that, for the Gaussian approximation to be valid, the coexistence, interior maxima must be located far away from the boundaries, so that the joint probabilities associated to all of extinction states are negligible.

The results of this analysis are summarized in Fig. 5. For low levels of stochasticity, the numerical coefficient of variation and the analytical approximation, Eq. (D.8), remain close to each other. For the corresponding values of *K*, the probability of coexistence is almost equal to one (as in the deterministic scenario). However, as *v* increases (i.e., the carrying capacity *K* decreases), the probability that one species becomes extinct grows sharply and, at the same time, the probability of coexistence starts declining. As the variability in population sizes augments, overlapping windows for the likelihood of progressive extinctions arise. The transition to situations where low demographic stochasticity operates is, therefore, abrupt.

**Figure 5:**
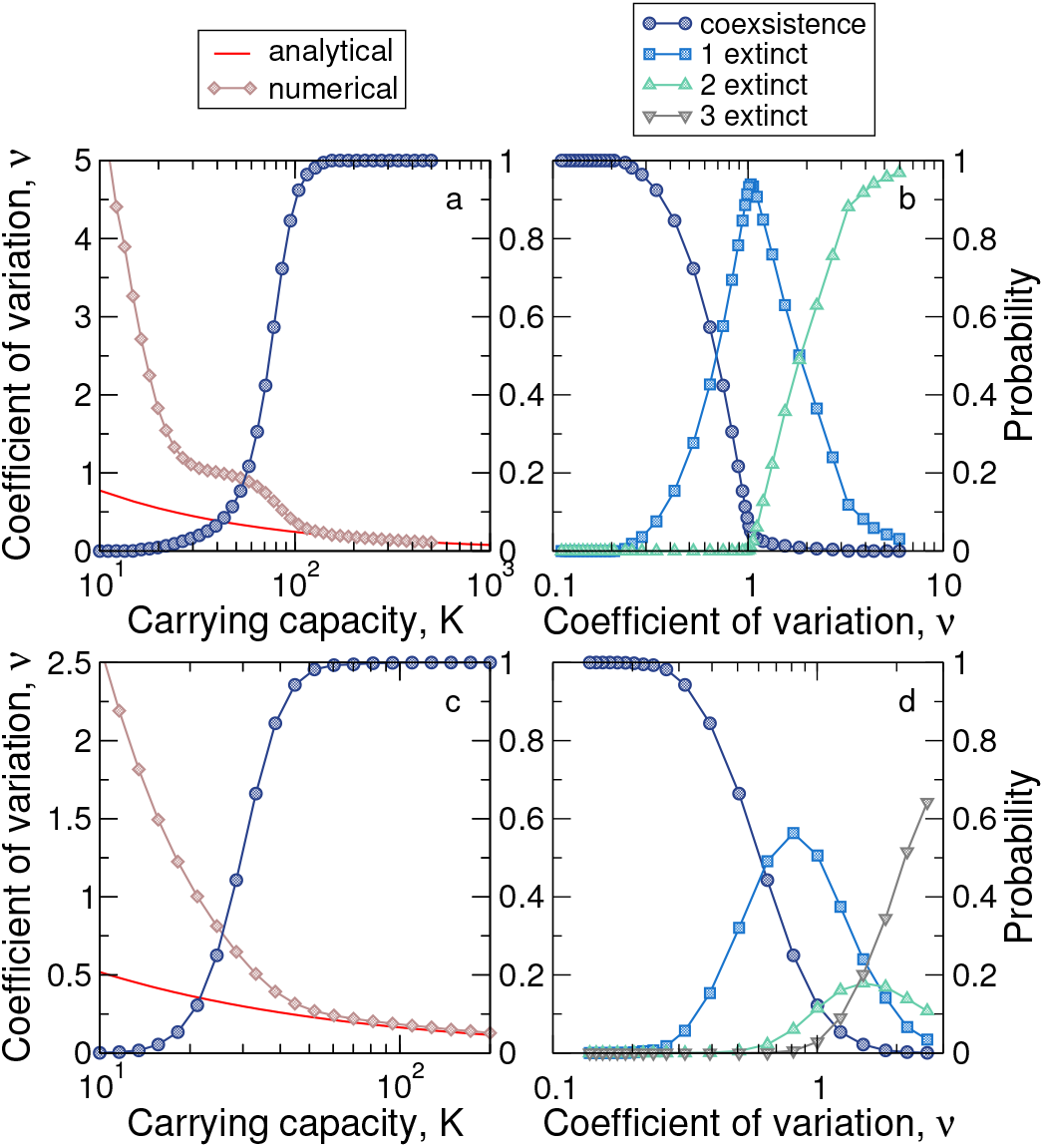
Cascade persistence for variable levels of demographic stochasticity. Panel (a) shows the coefficient of variation of population abundances, *v* = *σ_n_*/(*n*) (diamonds), as a function of increasing carrying capacity, *K*, for *S* = 2, *r*^+^ = 50, *r*^−^ = 25, *μ* = 0.1 and *ρ* = 0.5. For large *K*, the coefficient of variation tends to the analytical approximation (D.8) derived in Appendix D. Simultaneously, the probability of coexistence (alternative vertical axis, circles) becomes closer to 1. (b) Extinction cascade as a function of demographic stochasticity. At *v* ~ 0.2, the probability that one species goes extinct abruptly increases, and the coexistence probability starts declining. As *v* increases, higher-order extinctions become more likely. Panels (c)-(d) are equivalent to (a)-(b) but for *S* = 3, *r*^+^ = 10, *r*^−^ = 5, *μ* = 0.1 and *ρ* = 0.1. When the Gaussian approximation is valid, coexistence probability is almost equal to 1. After an abrupt decline, sequential extinctions occur for higher levels of demographic noise.

Note that the cascade obtained in Fig. 5b,d as a function of noise can be immediately translated into a cascade in carrying capacity (for fixed *ρ*). This reinforces our conclusion, since the extinction cascade phenomenon also occurs when other model parameter (*K*) varies. It is presumably the relative balance between *ρ* and *K* that determines the subspace of the parameter space for which extinctions start appearing.

### 3.5. Evaluating the role of environmental stochasticity

In order to test the robustness of our main result against different sources of noise, we have replaced demographic stochasticity by environmental stochasticity. We have introduced variability in model parameters so that, to keep the scenario as simple as possible, the competitive overlap *ρ* in Eq. (1) is replaced by *ρ* + *ξ*(*t)*, where *ξ*(*t*) stands for a noise term with zero mean. The deterministic dynamics transforms into a Langevin equation with multiplicative noise,

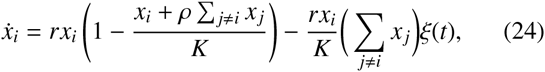

which can be rewritten as

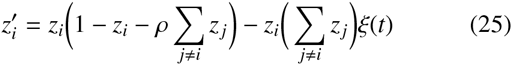

by re-scaling species densities as *z_i_* = *x_i_*/*K* and time as *t′* = *rt* (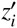 stands for the derivative with respect to the scaled time *t′*). We choose the noise as *ξ*(*t*) = *ρκη*(*t*), where *η*(*t*) is a Brownian motion. The noise has been scaled by *ρ* in order to avoid that the overall competitive strength, *ρ* + *ξ*(*t*), becomes negative.

Fig. 6 shows the persistence of the extinction cascade when only environmental stochasticity is considered. This points to the robustness of our results: as for demographic stochasticity, environmental stochasticity also alters the predictions of the deterministic dynamics. There are, however, qualitative differences between the predictions yielded by the model when demographic or environmental stochasticity come into play. First, the range in competition on which the cascade takes place is wider for demographic noise. It seems that, in the presence of environmental noise, the range of the cascade can increase moderately when a larger number of species are to be packed (Fig. 6b). More importantly, a second difference arises: no full extinction is possible in the case of environmental noise. This is a peculiarity, not altered by the noise, of a generic competitive, Lotka-Volterra dynamics (cf. Eq. (A.2) in Appendix A), for which it can be easily shown that the full extinction equilibrium (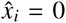, *i* = 1,…, *S*) is unstable. The Langevin equation, therefore, can not reproduce configurations associated to the full extinction of the community, contrary to what is observed for demographic stochasticity (Fig. 4b).

**Figure 6:**
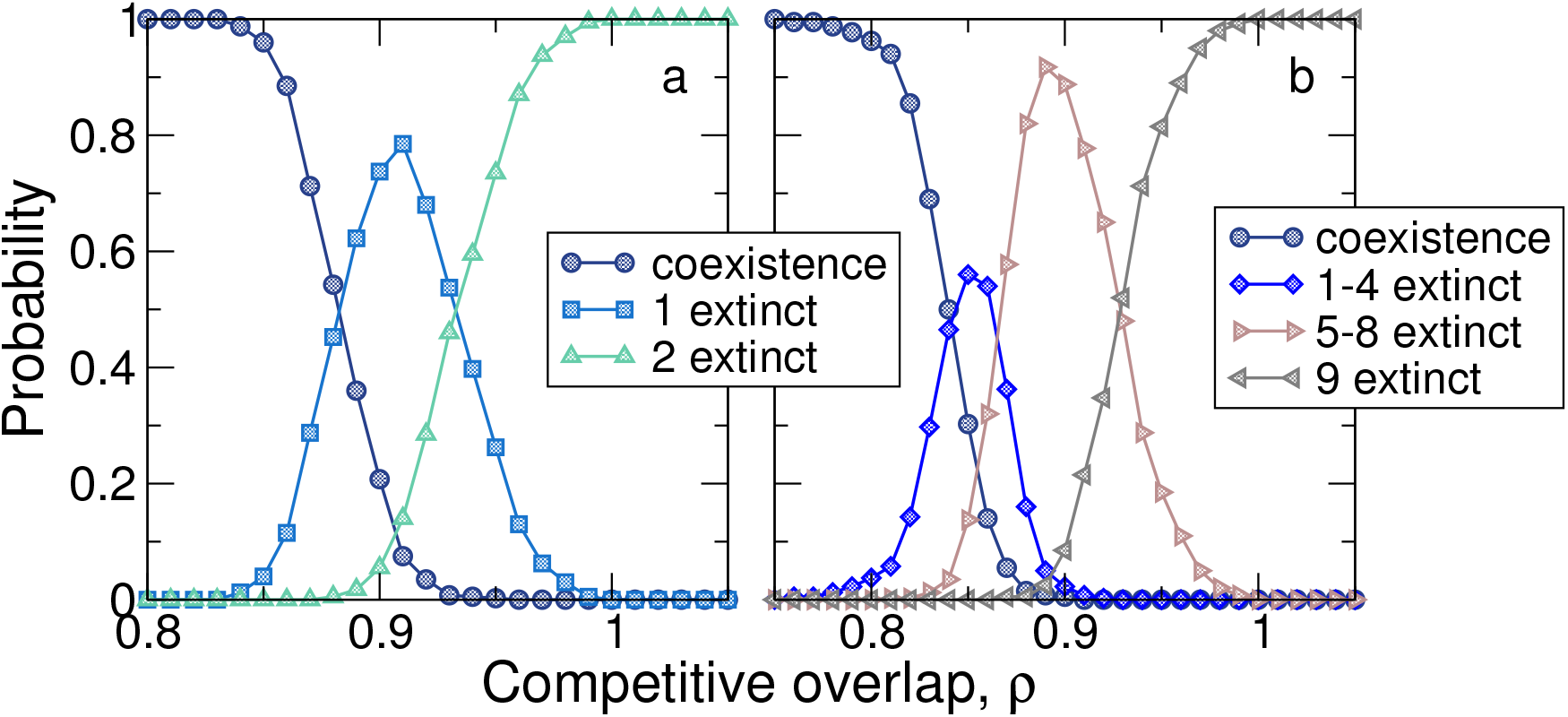
Cascade of extinctions in the case of environmental stochasticity. The Langevin equation (25) has been numerically integrated for (a) *S* = 3 and (b) *S* = 10 species. At the end of the simulated time span, species i is regarded as extinct if the corresponding density *z_i_* is at least 1% smaller than the maximum density *z_m_* = max{*z*_1_,…, *z_S_*}. Probabilities are calculated here by averaging over 400 realizations starting from random initial conditions. Here we take *κ* = 0.5 in the definition of the multiplicative noise *ξ*. To ease visualization, in panel (b) we have aggregated together the probabilities from one to four extinctions, as well as the probabilities of observing from five to eight extinctions. Consistently with the deterministic model, the full extinction state is never observed even in the presence of noise.

### 3.6. Relaxing the species symmetry assumption

In this section we test the robustness of our results by relaxing the assumption of species symmetry in model parameters. It can be argued that the effect of demographic stochasticity simply consists of breaking the symmetry between species. In a generic, non-symmetric, deterministic scenario, one could expect progressive extinctions even in the absence of ecological drift. Therefore, the role of stochasticity would be simply to re-establish a deterministic scenario where one-by-one species extinctions occur. In this subsection we discuss the implications of relaxing the symmetry assumption to determine the true role of demographic stochasticity in a generic case.

In order to address these questions, here we consider two examples of fully non-symmetric, three-species competitive dynamics,

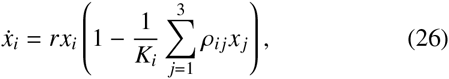

where carrying capacities and interspecific competitive strengths are species-dependent. We set, without loss of generality, off-diagonal competition values as *ρ*_12_ = *ρ*_21_ = *ρ*, *ρ*_13_ = *ρ*_31_ = *ρ* + *δ*_1_, *ρ*_23_ = *ρ*_32_ = *ρ* + *δ*_2_, and *ρ_ii_* = 1 for *i* = 1,2,3.

In the first example we choose *δ*_1_ = 0.1, *δ*_2_ = 0.05, *K*_1_ = 40, *K*_2_ = 16, and *K*_3_ = 20. After analyzing the stability of all the equilibrium points of the deterministic system (see details in Appendix E) we find that, although being fully non-symmetric, this model predicts a two-species, grouped extinction when the threshold *ρ* = 0.4 is crossed over (Fig. 7a). For *ρ* > 0.4, only equilibria with a single extant species are stable, whereas no other scenario but three-species coexistence is stable for 0 ≤ *ρ* < 0.4. By continuity of the eigenvalues of the Jacobian matrix on model parameters, there are multiple non-symmetric, deterministic scenarios that exhibit the transition from full coexistence to a single-extant-species state. Therefore, grouped extinctions are not a peculiarity of the fully symmetric scenario.

**Figure 7:**
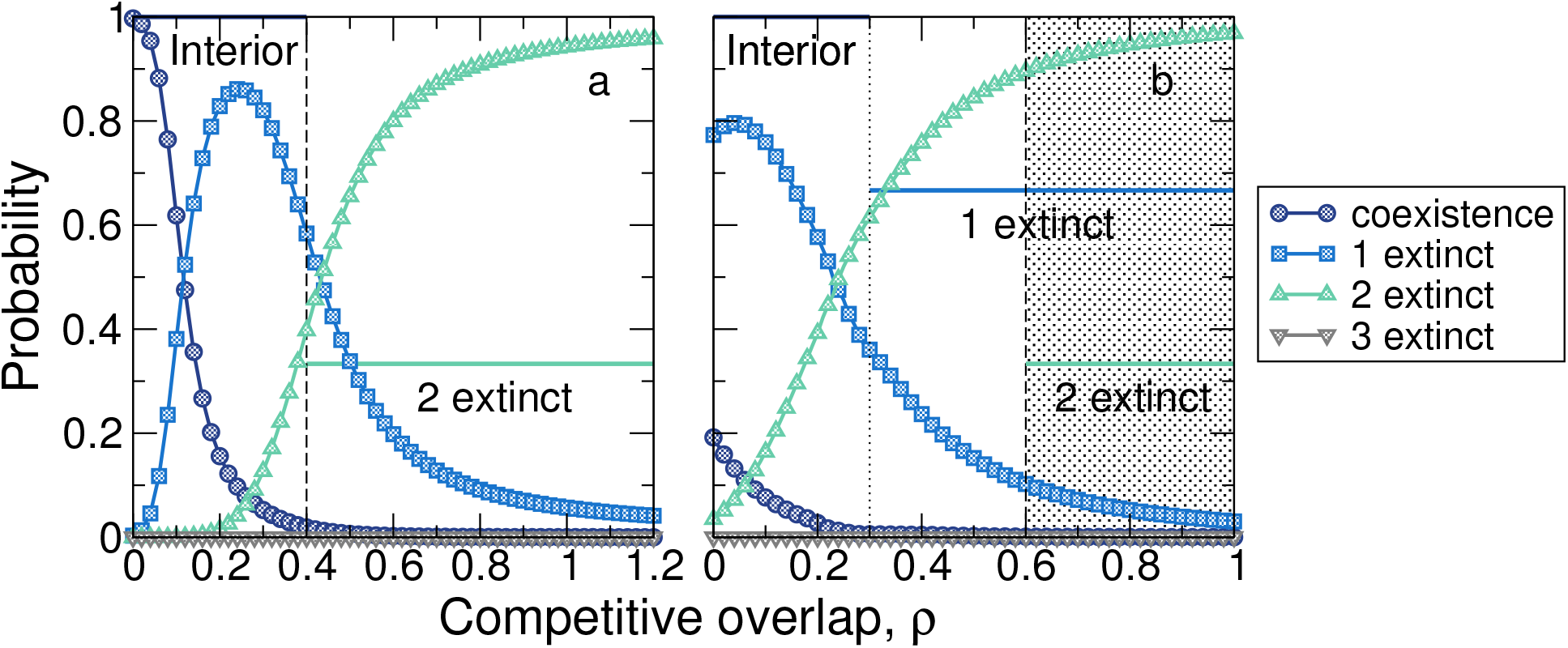
Non-symmetric scenario described in Sec. 3.6. In both panels, *r*^+^ = 5.1, *r*^−^ = 0.1 and *μ* = 0.1. (a) In this case, the deterministic model predicts a multiple extinction of two species when the threshold *ρ* = 0.4 is crossed over (the ranges where the deterministic model yields stability are marked with horizontal lines). Remaining parameters are *K*_1_ = 40, *K*_2_ = 16, *K*_3_ = 20, *δ*_1_ = 0.1 and *δ*_2_ = 0.05. The stochastic scenario, however, yields a sequential cascade for which extinctions are spread out along the axis of competitive overlap (symbols). (b) Here carrying capacities are uniform (*K*_1_ = *K*_2_ = *K*_3_ = 15) and *δ*_1_ = 0.4 and *δ*_2_ = 0.3. The stability analysis of the deterministic model implies ranges where multiple stable equilibria co-occur (shadowed area). The extinction sequence is not fully determined, and has to be compared with the stochastic prediction.

In the second example (Fig. 7b), *δ*_1_ = 0.4, *δ*_2_ = 0.3 and *K*_1_ = *K*_2_ = *K*_3_ = 15. The complete stability analysis draws the conclusion that non-symmetric models yield to multistability. In Appendix E it is shown that, for *ρ* < 0.3, only the interior equilibrium point is asymptotically stable; for 0.3 < *ρ* < 0.7, only the boundary point (15/(1 + *ρ*), 15/(1 + *ρ*), 0) is stable, but for 0.7 < *ρ* < 1, the former boundary point remains stable as well as the single-species extinction equilibrium (0,0,15) (see also Fig. 7b). For *ρ* > 1, however, the three boundary equilibria with a single, extant species are the only ones that remain stable. As a consequence, there is a range in competition (0.7 < *ρ* < 1) where configurations formed by a single extant species or by two coexisting species co-occur. Depending on the initial condition, the dynamics can end up in one of the two attractors. The basin of attraction of each equilibrium point will determine how frequently one or two species go extinct when competition surpasses the value *ρ* = 0.7. This analysis is out of the scope of this contribution, though. Importantly, the extinction sequence in non-symmetric scenarios can depend on initial conditions and is not fully determined in principle.

Multiple extinctions are not precluded in general even when the symmetry between species is broken, as the first example shows. Multistability ranges could also lead to grouped extinctions in deterministic scenarios. In a Lotka-Volterra community model with *S* species there are 2^*S*^ attractors. Increasing complexity would likely lead to additional overlapping regions in competition where multiple stable attractors co-occur and species can decline together. Moreover, the deterministic extinction can be ambiguous. It seems difficult to establish the conditions under which a general non-symmetric model will produce a well-defined extinction sequence. The variability introduced by idiosyncratic, species-dependent carrying capacities, growth rates, or intra- and interspecific strengths, may cause the extinction sequence to be analytically unpredictable for species-rich communities.

Remarkably, the extinction sequence predicted by non-symmetric, deterministic models has nothing to do with the cascade observed when demographic stochasticity comes into play. We have calculated the probabilities of coexistence and one-, two- or three-species extinctions for both stochastic parametrizations of the two non-symmetric models considered in this subsection. To aggregate joint probabilities, we obtained numerically the critical points by spline interpolation of the exact joint distribution. Now the six saddle points have asymmetric entries, but three of them exhibit a coordinate close to the boundary, and the remaining three saddle points present two coordinates near the axes. Thus, the partition of the configuration space is conceptually equivalent to that of Fig. 3. Not surprisingly, as Fig. 7 evidences, the stochastic model predicts a sequential cascade of extinctions. The threshold at which extinctions start occurring displaces towards smaller values of p, leading to narrow ranges of effective stochastic coexistence. The most likely state in the presence of stochasticity is not necessarily the same as in a deterministic scenario, the predictions being utterly different in terms of the extinction sequence. Therefore, regardless of the inherent lack of symmetries that deterministic dynamics may have, demographic stochasticity can influence significantly the way in which extinctions take place.

## 4. Discussion

In this contribution, we have analyzed the extinction phenomenon for a symmetric, Lotka-Volterra competitive system formed by *S* species. In particular, we have focused on the differences between the deterministic system and its stochastic counterpart. Our main result is related to the way in which species extinction proceeds: on the one hand, in the deterministic system, *S* – 1 species are driven to extinction at the very point where competitive exclusion starts to operate. On the other hand, we have shown that the overall probabilities of coexistence and one-, two-, or three-species extinction alternate sequentially as the most likely states when competitive overlap increases. Therefore, stochasticity is responsible of a progressive sequence of species extinction, a phenomenon that is absent in the deterministic system. Our analyses are based on a birth-death-immigration stochastic dynamics that was analyzed in deep by Haegeman and Loreau (2011) and later used by Capitán et al. (2015) to unveil the existence of a more restrictive threshold in competition when demographic stochasticity is explicitly considered. In addition to lowering the threshold in competition at which extinctions begin to occur, ecological drift also changes drastically the way those extinctions take place. In order to evidence this difference, we have developed convenient analytical approximations to the critical points of the joint probability for *S* = 2 and *S* = 3 potential species, which were used to divide the configuration space of the stochastic process to yield aggregated probabilities associated to coexistence and to the corresponding configurations with one or more extinct species. These probabilities reveal the stochastic extinction cascade.

We have shown that different community configurations alternate within certain ranges of competition: coexistence and one-species extinction alternate for small *ρ*, whereas one-species and two-species extinction most likely interchange among each other for intermediate values of *ρ*. The presence of multiple modes in the joint probability distribution is akin to the presence of multistability in a deterministic system. However, the symmetric, deterministic model is characterized by the absence of multiple stable states for *ρ* < 1. Hence the extinction cascade described in this work is an entirely new effect caused only by ecological drift. In addition, we have evidenced that the transition to the deterministic model is sharp when the intensity of demographic stochasticity tends to zero. Moreover, we have proven that environmental stochasticity also leads to a cascade of extinctions, although the ranges in competition where extinctions take place are smaller and, more importantly, the full extinction of the community is not possible in this case.

In this work, we have implemented two types of stochasticity: demographic noise (ecological drift) and environmental stochasticity. These are two typical sources of noise that represent, respectively: (i) the variability in discrete population numbers as a consequence of stochastic births and deaths, or (ii) the stochastic variability in model parameters that can be ascribed to changing environmental conditions. Although they are very different implementations of noise, the main result of this manuscript (the stochastic cascade of extinctions) is common to both of them, with some qualitative differences. In the case of demographic stochasticity, we do not impose a particular form for the noise distribution since it directly emerges from the inherent stochastic dynamics of discrete populations whose individuals undergo a number of elementary processes (in principle, the noise distribution would follow from the master equation). Other ways to implement demographic noise have been discussed in the literature (Bonachela et al., 2012), and they would plausibly lead to mechanisms similar to those found here.

It can be argued that the role of stochasticity in the fully symmetric system reduces to break species symmetry and yield to progressive species extinctions, a scenario that can arise in non-symmetric, deterministic approaches. We have illustrated with examples that grouped extinctions are not exclusive of a fully symmetric situation. Moreover, predicting the extinction sequence in non-symmetric, deterministic cases is difficult because multiple stable equilibria can co-occur in ranges of competition. Besides, although the extinction sequence were completely determined, the cascade in the presence of stochasticity can be totally different from that predicted by the deterministic model. We believe that the examples analyzed in this contribution show up the key role of stochasticity in community assembly.

It would be interesting to empirically test the stochastic extinction cascade phenomenon. In principle, a plausible way to conduct the experiment would involve simple protist microcosms where species compete for a shared resource (see, for example, Violle et al. (2010) and references therein). Lowering the amount of resource could be associated to a decrease in the carrying capacity, and we have shown that the extinction cascade mechanism is expected to arise as long as the carrying capacity (resource availability) is reduced, see Fig. 5. For the experiment to reproduce model settings, an individual (immigrant) coming from the species pool should be inserted in the experimental community at certain times. From a time series listing species identities at certain sampling times, one could estimate the probabilities for coexistence and for the extinction of one or more species, and test whether extinctions in empirical systems tend to proceed sequentially or not.

Our work has two important implications: first, we have developed analytical approximations for conditional probabilities in the cases *S* = 2 and *S* = 3. As we have shown, these functions work well at least around the critical points of the joint probability. Presumably, the techniques proposed here might be extended to approximate the joint probability distribution itself. This approach, however, has to be performed carefully. The simplest way to approximate the three-species joint distribution according to our methodology is to set *P*(*n*_1_, *n*_2_, *n*_3_) ≈ *T*(*n*_1_|*n*_2_, *n*_3_)*T*(*n*_2_|*n*_3_)*P*(*n*_3_), where *P*(*n*_3_) is the marginal, one-species probability distribution, which can be expressed analytically in terms of hypergeometric functions (Haegeman and Loreau, 2011). Apparently, the approximated distribution lacks of an important property of the exact joint distribution: it is not conserved under cyclic permutations of its arguments. Therefore, it is necessary to devise appropriate combinations of conditional probabilities that preserve the symmetry of the joint distribution under cyclic permutations of its arguments. This research direction, together with a generalization for communities formed by more than three species, could be worth pursuing and compared with other approximations to the joint probability, such as those developed by Haegeman and Loreau (2011).

The second implication of this contribution is that we remark the importance of explicitly considering ecological drift in theoretical frameworks in community ecology. Natural processes are intrinsically stochastic, because changes in population numbers are discrete, so ecological communities are more reliably modeled using stochastic community models, even at regimes where their deterministic limits are not expected to fail. In certain situations, the use of deterministic dynamics in community ecology could lead to utterly different predictions, as we have shown. Traditionally ecology has relied on these kind of models, and currently considerable theoretical progress is made on the basis of deterministic approaches. However, ecological drift may play a determinant role that could not be captured by deterministic formulations. We hope that the present work can inspire new contributions in the future that highlight the distinct role of ecological drift in species community models.

## 5. Acknowledgements

We thank the constructive criticisms and comments of the Editor and two anonymous reviewers, who helped to improve significantly the original manuscript. This work was funded by project SITES (CGL2012-39964, DA, JAC) and the Ramon y Cajal Fellowship program (DA). JAC also acknowledges partial financial support from the Department of Applied Mathematics (Technical University of Madrid).

## Appendix A. Deterministic competitive exclusion

Gause’s competitive exclusion principle is usually stated as “two species competing for a single resource cannot coexist”. Strictly speaking, the competitive exclusion principle was first formulated by Volterra (1926) as a mathematical proposition. Albeit a mathematical proof of the principle, based on dynamical systems theory, can be found in the book by Hofbauer and Sigmund (1998), and is essentially the same as that of Volterra (1926) original paper, we here provide a purely algebraic alternative demonstration (see Roughgarden (1979) as well).

Assume that the densities of S species living on R resources vary in time according to the dynamics

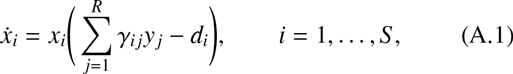

where Γ = (*γ_ij_*) is a *S* × *R* matrix with non-negative entries, *y_j_*, *j* = 1,…, *R*, are amounts of *R* resources, which are also assumed to depend on species densities, and *d_i_, i* = 1,…, *S* are the rates of decline of species when all resources are zero. We now show that if *S* > *R* and species densities reach a well-defined steady state, then in the long run *S* – *R* species will go extinct until the number of species equals the number of resources.

Imposing the condition 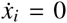 for large *t* and assuming that all species densities are positive, we get the non-homogeneous linear system Γ**y** = **d**, with 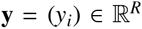, and 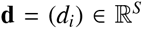. Given that matrix Γ has *S* rows and only *R* < *S* columns, the system will be incompatible except for some specific vectors **d** belonging to the image Im(Γ) of matrix Γ. As a consequence, to find an equilibrium solution some species densities must go to zero. For those displaced species, the corresponding rows of matrix Γ can be removed until rank(Γ) rows remain. If the rank equals the number *R* of resources, the system turns out to be compatible and determinate, and *R* species will stably coexist. Note that it is the rank of matrix Γ rather than the number of resources itself that induces competitive exclusion.

Therefore, competition for shared resources imposes a limit to the maximum number of species that can stably coexist. This result has a counterpart in the stability of Lotka-Volterra equations derived from the MacArthur’s consumer-resource model (MacArthur, 1970; Chesson, 1990), i.e., the particular case of Eq. (A.1) in which per-capita resource growth rates 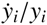 depend linearly on population densities. In the limit of fast resource variation, the dynamics (A.1) takes the Lotka-Volterra form (Chesson, 1990),

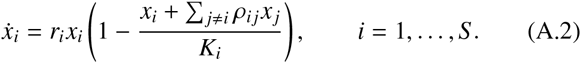

Here *r_i_* is interpreted as the intrinsic growth rate of species *i, K_i_* as a carrying capacity, and *ρ_ij_* measures interspecific competition strength between species *i* and *j* relative to intraspecific competition. As shown by Chesson (1990), if *S* = *R* = rank(Γ), then exists a unique solution of the system 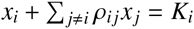, *i* = 1,…, *S*. The resulting equilibrium point will be interior if every species density remains strictly positive. Moreover, Chesson (1990) demonstrated that if there exists a unique interior equilibrium point for the dynamics (A.2), it will be globally stable if and only if *S* = *R* = rank(Γ). Therefore, if competitive exclusion does not operate, a unique globally stable equilibrium point is reached and, conversely, if the Lotka-Volterra equations present an interior, globally stable equilibrium point, any positive initial condition will make the dynamics (A.2) converge to the equilibrium point. This automatically ensures that none of the *S* species is driven to extinction by competitive exclusion (MacArthur, 1970; Takeuchi, 1996).

## Appendix B. Stability of the symmetric, deterministic model for *ρ* ≥ 1

In this Appendix we perform a stability analysis of the equilibrium points of the symmetric, deterministic dynamics in the competition regime *ρ* ≥ 1. As we mentioned before, when *ρ* < 1 it can be shown that the interior equilibrium point is globally stable (Hofbauer and Sigmund, 1998; Capitán et al., 2015), all boundary equilibria being unstable.

We start by analyzing stability for *ρ* = 1. In this case, the system is also stable and the initial condition determines the attractor which the dynamics converges to. Any equilibrium point is such that its densities satisfy

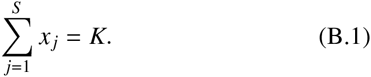

Let 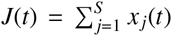. Then, by summing up the equations of the system (1) for *ρ* = 1 we get

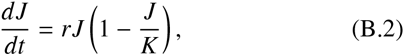

which can be integrated and yields

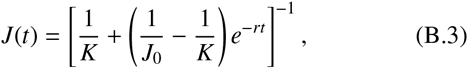

*J*_0_ being the initial condition for *J*(*t*), 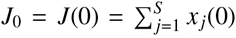. Therefore, the dynamics of each species turns out to be decoupled,

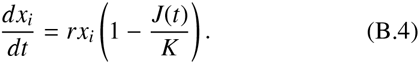

The equilibrium point to which (B.4) converges is determined by the initial condition vector **x_0_** = (*x*_1_(0),…, *x_S_* (0)). For two distinct species *i* and *j*, (B.4) implies that 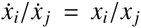. Integration yields *x_i_*(*t*)/*x_j_*(*t*) = *x_i_*(0)/*x_j_*(0), which means that the proportions of population densities are conserved along the dynamics, whose orbits are reduced to straight lines starting from the initial condition **x**_0_ along the direction determined by the vector **x**_0_. Therefore, the final equilibrium point is given by the intersection of the hyperplane 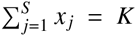 and the line that links the initial condition point, **x_0_**, and the origin. Any of these (infinite) equilibrium points will be stable, provided that the initial densities satisfy *x_i_*(0) ≥ 0 for all *i*.

The case *ρ* > 1 leads to competitive exclusion. We analyze the asymptotic stability of the 2^*S*^ equilibrium points with positive or zero densities. Without loss of generality, for 0 ≤ *n* ≤ *S* any equilibrium point will be of the form

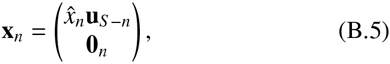

where subscript *n* indicates that *n* densities are strictly equal to 0, and **u**_*B*_ = (1,…, 1)^*T*^ is a vector with n entries equal to 1. Any equilibrium with *n* zero densities can be written as a permutation of (B.5), without altering the subsequent stability analysis. In addition, the non-zero entries of **x**_*n*_ are the solutions of the system

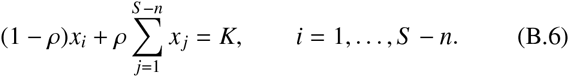

This system admits a single solution for which the *S* – *n* species have equal densities, 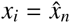, where

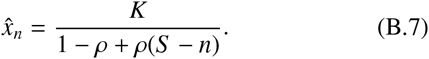

Our demonstration reduces to show that, if *ρ* > 1, all the nontrivial equilibria act as repellors except for *n* = 1, i.e., when only a single species survives. To this purpose we evaluate the eigenvalue spectra of the Jacobian matrix of the system (1). The Jacobian matrix J can be expressed in a block form as

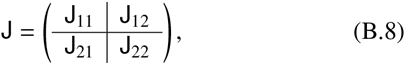

where

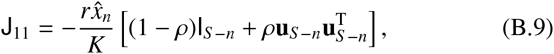

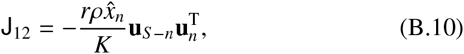

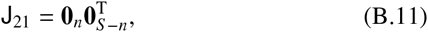

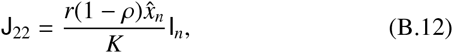

and I_*n*_ is the *n* × *n* identity matrix and **0**_*n*_ = (0,…, 0)^T^ is the zero vector with n entries. Without loss of generality, eigenvectors can be written as

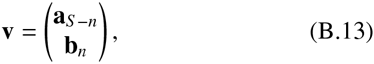

where **a**_*S–n*_ and **b**_*n*_ are column vectors with dimensions *S* – *n* and *n*, respectively. First let us assume that **b**_*B*_ = **0**_*n*_. The spectral problem reduces to

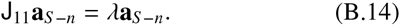

Two different solutions arise: if 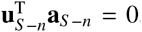, i.e., **a**_*S–n*_ is orthogonal to **u**_*S–n*_, then the eigenvalue is

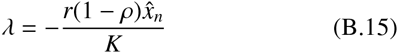

with algebraic multiplicity *S* – *n* – 1. However, if **a**_*S–n*_ ∝ **u**_*S–n*_ we find *λ* = –*r* as an eigenvalue with algebraic multiplicity equal to 1.

On the other hand, if **b**_*n*_ ≠ **0**_*n*_ we have to solve

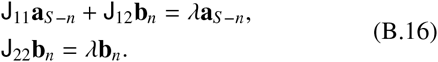

Since J_22_ is proportional to I_*n*_, we find

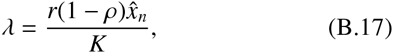

with *n* eigenvectors 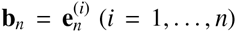, 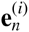 being the *i*-th vector of the canonical basis of 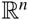. The algebraic multiplicity associated to (B.17) is equal to n. Substituting these results into the first equation of (B.16) yields

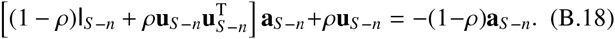

Given the structure of the matrices involved, we look for a solution of the form **a**_*S–B*_ = *α***u**_*S–n*_, which implies a non-trivial value *α* = –*ρ*/[2(1 – *ρ*) + *ρS*].

Two eigenvalues determine the asymptotic stability of all the equilibrium points: 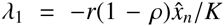 and 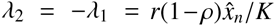 (the third eigenvalue, *λ*_3_ = –*r*, is always negative). If *ρ* > 1 and 0 ≤ *n* < *S*–1, *λ*_1_ > 0 and remains as an eigenvalue—recall that its multiplicity is *S* – *n* – 1 > 0. Therefore, any equilibrium with less than *S* – 1 extinct species is asymptotically unstable—including the interior coexistence equilibrium. However, when only one species survives (*n* = *S* – 1), *λ*_1_ is not an eigenvalue anymore and the two other eigenvalues remain: 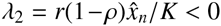 (with multiplicity *S*–1) and *λ*_3_ = –*r* < 0 (with multiplicity 1). Thus, only the S boundary equilibria with a single extant species are asymptotically stable.

Since the trivial equilibrium point (associated to complete extinction) is obviously unstable, we deduce that any orbit starting from an interior initial condition will be repelled if it gets close to any equilibrium, except when the equilibrium point is formed by a single extant species. If the orbit enters the basin of attraction of any of those *S* equilibria, it will end up in it asymptotically. This implies the extinction at a time of *S* – 1 species if *ρ* > 1.

## Appendix C. Numerical calculation of the steady-state probability distribution

In order to compute numerically the stationary joint distribution, we limit the infinite configuration space to the set Ξ {0,1,…, *n_max_*}^*S*^ by choosing *n*_max_ large enough so that the probability of finding a population number equal to *n*_max_ is negligible (we choose *n*_max_ as the integer part of 2K, which fulfills the requirement).

The stationary distribution is obtained by solving the embedded Markov chain associated to the continuous-time Markov process (Karlin and Taylor, 1975). The transition matrix of the embedded Markov chain is defined by the transition probabilities

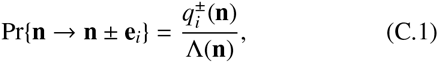

where 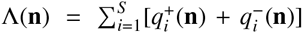. The remaining transitions have zero probability. Elementary events (overall births and deaths) take place after exponential times, so that the time lapsed to the next event is drawn from a random variable *τ* with cumulative distribution Pr(τ ≤ *t*) = 1 – *e*^−Λ(**n**)*t*^. Once the steady-state distribution ***φ*** = (*φ*(**n**)) of the embedded Markov chain—i.e., the left-eigenvector of the transition matrix with eigenvalue 1—has been determined, according to the mean time spent by the process at state **n**, the probability of finding the continuous-time Markov process at state **n** turns out to be (Cinlar, 1975)

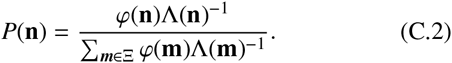

## Appendix D. Coefficient of variation of population abundances in the limit of large carrying capacity

In this section we derive an analytical expression for the coefficient of variation of population abundances in the small stochasticity limit. In the case of demographic stochasticity, low variability levels can be obtained for large population sizes or, equivalently, in the limit of large carrying capacity. We build on the Gaussian approximation for the joint probability distribution, which is valid in the limit *K* ≫ 1 since the probability of extinction configurations is expected to be negligible, and can be fully calculated for a generic community of size *S*.

The Gaussian approximation can be obtained as the solution to the Fokker-Planck equation deduced from the master equation (2). We do not reproduce its derivation here; it can be found at the Supplemental Information of Capitán et al. (2015). Under this approximation, the joint probability distribution is expressed as

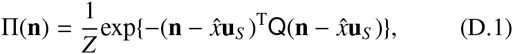

where *Z* is an appropriate normalization factor,

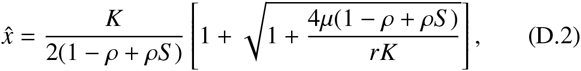

and the covariance matrix Q^-1^ is given by

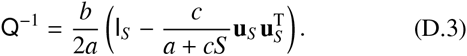

In terms of model parameters, 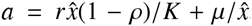, 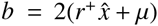, and 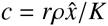. In the large carrying capacity limit, the average population abundance is expressed through a series expansion on powers of *K* as

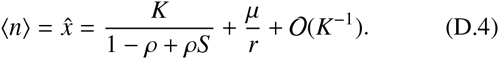

Similarly, series expansions give

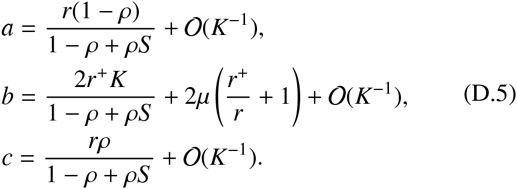

Inserting these expressions into (D.3) yields, up to order *K*^0^, the approximation

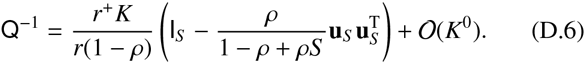

Therefore the standard deviation of population abundance, *σ_n_*, can be obtained as the square root of diagonal elements of matrix Q^-1^,

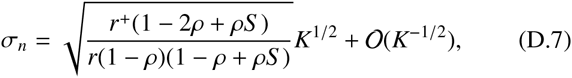

which (not surprisingly) scales with *K* as *K*^1/2^. Finally, the coefficient of variation of population abundances is expressed, in the limit *K* ≫ 1, as

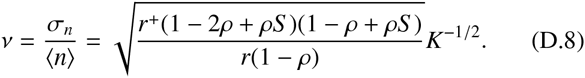

Strictly speaking, the deterministic scenario (*v* = 0) is only achieved in the limit *K* → ∞. However, low stochasticity regimes can be assessed using Eq. (D.8): if the actual coefficient of variation is close to that yielded by the Gaussian approximation, both of which are small for large *K*, then extinction configurations are precluded and the variability of populations with respect to the mean value is small. We adopt, as a practical definition for low stochasticity, the parameter combinations for which the actual coefficient of variation is close to the approximation given by Eq. (D.8).

## Appendix E. Stability analysis for two deterministic models with non-symmetric competition

Here we analyze the stability of the equilibrium points of the two non-symmetric, three-species competitive dynamics of the form (26) considered in the main text, for which the interaction matrix is written as

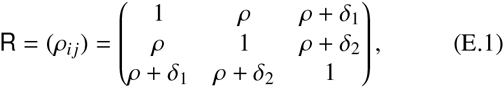

*δ*_1_ and *δ*_2_ being two positive numbers. The sign of equilibrium densities and the corresponding eigenvalues of the Jacobian matrix determine the ranges of ρ for which the system is stable. We impose the condition *ρ* ≥ 0 for all interaction coefficients to remain positive. Since the growth rate *r* > 0, in both cases the full extinction equilibrium 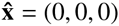 is unstable.

In the first example, *δ*_1_ = 0.1 and *δ*_2_ = 0.05. Species carrying capacities are non-uniform: *K*_1_ = 40, *K*_2_ = 16, and *K*_3_ = 20. Although the expressions are too cumbersome to be reproduced here, it can be shown that the interior equilibrium point 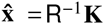, with **K** = (*K*_1_, *K*_2_, *K*_3_)^T^, has three positive densities and is asymptotically stable if and only if 0 ≤ *ρ* ≤ 0.4.

We now consider all boundary equilibria. In what follows the eigenvalues *λ* of the stability (Jacobian) matrix are expressed as *λ* = *λ′r*, i.e., they are scaled by the growth rate *r* > 0:

a. The first and second coordinates of

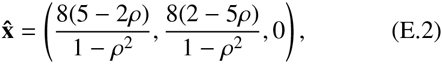

are positive if and only if 0 ≤ *ρ* ≤ 0.4 or *ρ* > 2.5. The scaled eigenvalues *λ′* are given by

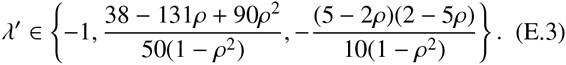 The condition 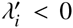 for *i* = 1,2,3 yields 1.055 < *ρ* < 2.5. Therefore, this point is never feasible and stable at the same time.
b. For

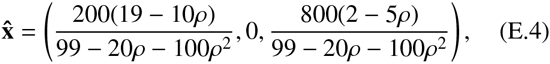

we require positivity for the first and third entries, which yields 0 ≤ *ρ* < 0.4 or *ρ* > 1.9. On the other hand, the eigenvalues in this case are

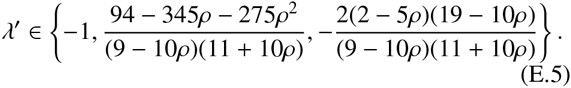 Stability implies 0.9 < *ρ* < 1.9, which is incompatible with the feasibility condition.
c. The third equilibrium with a single extinct species is

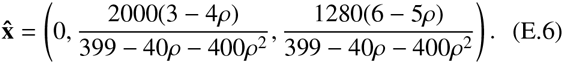 This point is feasible if and only if 0 ≤ *ρ* < 0.75 or *ρ* > 1.2. The (scaled) eigenvalues of the Jacobian matrix are

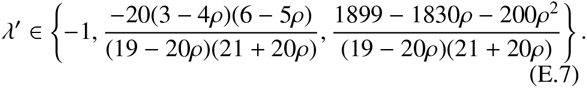 Since the system of inequalities 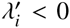 (*i* = 1,2,3) turns out to be incompatible, this point is unstable for all values of *ρ*.
d. 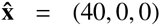: the scaled eigenvalues are {–1, (2 – 5p)/2,2(2 – 5*ρ*)/5}. This point is stable for *ρ* > 0.4.
e. 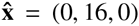: *λ′* ∈ {–1, (5 – 2*ρ*)/5, 4(6 – 5*ρ*)/25}. The equilibrium point is stable if and only if *ρ* > 2.5.
f. 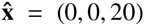: *λ′* ∈ {–1, (19 – 10*ρ*)/20, 5(3 – 4*ρ*)/16}. Asymptotic stability is achieved for *ρ* > 1.9.

As a result, in the range 0 ≤ *ρ* < 0.4, the only stable point is the coexistence equilibrium. However, for *ρ* > 0.4, only two-extinct species equilibria remain asymptotically stable. Since the eigenvalues are continuous functions of model parameters, close to this example we can find multiple non-symmetric systems that exhibit a grouped, two-species extinction as *ρ* increases.

The second example shows that multiple stable equilibria can co-occur when interactions are chosen non-symmetrically. In this case, we have taken *δ*_1_ = 0.4, *δ*_2_ = 0.3 and *K*_1_ = *K*_2_ = *K*_3_ = 15. The densities of the interior equilibrium point are expressed as

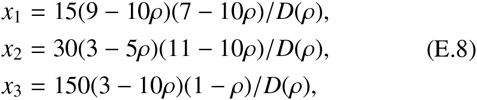

where *D*(*ρ*) = 75 – 116*ρ* – 160*ρ*^2^ + 200*ρ*^3^. The feasibility analysis of the equilibrium point yields, for *ρ* ≥ 0, the ranges 0 ≤ *ρ* < 0.3 or 0.7 < *ρ* < 0.9 or *ρ* > 1.1. The eigenvalues of the stability matrix can be fully calculated, although their expressions are too cumbersome to be reproduced here. It is easy to check that the three eigenvalues are negative if and only if 0 ≤ *ρ* < 0.3. Therefore, this equilibrium point is interior and asymptotically stable if and only if 0 ≤ *ρ* < 0.3.

We now summarize the stability analysis for boundary equilibria. All of them are feasible, so stability is only conditioned by the sign of eigenvalues (which are all real):

a. 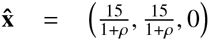: the scaled eigenvalues *λ′* are 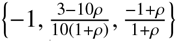. Therefore, this point is stable for 0.3 < *ρ* < 1.
b. 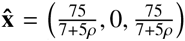: 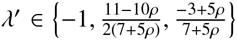. The stability conditions form an unfeasible problem, so this point turns out to be unstable for any *ρ*.
c. 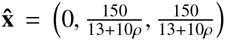: 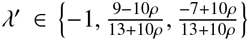. Again this point is unstable for all values of competitive overlap.
d. 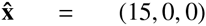: the scaled eigenvalues are 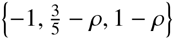. This point is stable for *ρ* > 1.
e. 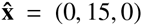: 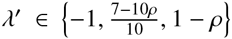. The equilibrium point is stable if and only if *ρ* > 1.
f. 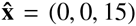: 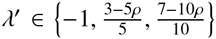. Stability is attained for *ρ* > 0.7.

Consequently, in the range 0.3 < *ρ* < 0.7 the only stable equilibrium point is 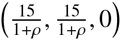. However, for 0.7 < *ρ* < 1 two stable equilibria co-occur: the former and a two-extinct species equilibrium, (0, 0, 15). Depending on initial conditions, the dynamics can lead to one of them or to the other. For *ρ* > 1, however, the three equilibria with a single extant species are the only ones that remain asymptotically stable.

This example shows how the cascade of extinctions in non-symmetric, deterministic models can be far from being determined due to the co-occurrence of multiple stable equilibria.

